# Rapid energy expenditure estimation for assisted and inclined loaded walking

**DOI:** 10.1101/401836

**Authors:** Patrick Slade, Rachel Troutman, Mykel J. Kochenderfer, Steven H. Collins, Scott L. Delp

## Abstract

**Background:** Estimating energy expenditure with indirect calorimetry requires expensive equipment and provides slow and noisy measurements. Rapid estimates using wearable sensors would enable techniques like optimizing assistive devices outside a lab. Existing methods correlate data from wearable sensors to measured energy expenditure without evaluating the accuracy of the estimated energy expenditure for activity conditions or subjects not included in the correlation process. Our goal is to assess data-driven models that are capable of rapidly estimating energy expenditure for new conditions and subjects.

**Methods:** We developed models that estimated energy expenditure from two datasets during walking conditions with (1) ankle exoskeleton assistance and (2) various loads and inclines. The estimation was portable and rapid, using input features that are possible to measure with wearable sensors and restricting the input data length to a single gait cycle or four second interval. The performance of the models was evaluated for three use cases. The first case estimated energy expenditure during walking conditions for subjects with some subject specific training data available. The second case estimated all conditions in the dataset for a new subject not included in the training data. The third case estimated new conditions for a new subject. The models also ordered the magnitude of energy expenditure across all conditions for a new subject.

**Results:** The average errors in energy expenditure estimation during assisted walking conditions were 4.4%, 8.0%, and 8.1% for the three use cases, respectively. The average errors in energy expenditure estimation during inclined and loaded walking conditions were 6.1%, 9.7%, and 11.7% for the three use cases. The models ordered the magnitude of energy expenditure with a maximum and average percentage of correctly ordered conditions of 56% and 43% for assisted walking and 85% and 55% for incline and loaded walking.

**Conclusions:** Our data-driven models determined the accuracy of energy expenditure estimation for three use cases. For experiments where the accuracy of a data-driven model is sufficient, standard indirect calorimetry can be replaced. The energy expenditure ordering could aid in selecting optimal assistance conditions. The models, code, and datasets are provided for reproduction and extension of our results.

## Background

The U.S. has an estimated 20 million people with ambulatory disabilities due to age, injury, disease, amputation, or congenital conditions [1]. These disabilities often result in less efficient gait patterns. Energy expenditure, or metabolic cost, is an important metric for understanding the level of effort required during motion [2,3]. Reducing this level of effort with gait retraining or assistance requires energy expenditure estimates to adapt the assistance to each user. Optimizing gait parameters and device assistance with “human-in-the-loop” methods has significantly reduced energy expenditure for steady state walking [4, 5, 6, 7]. These methods rely on indirect calorimetry to estimate energy expenditure, which is expensive and provides noisy measurements after each breath [8]. The estimation noise from variation between breaths is accounted for by averaging over a few minutes of breath measurements during steady state motions.

Methods for energy expenditure estimation span statistical and data-driven approaches as well as techniques that model the underlying mechanics and biological processes. Initial statistical methods of estimating energy expenditure fit linear equations from indirect calorimetry measurements of subjects moving at different speeds, inclines, or with additional loads [9,10,11]. Models based on walking mechanics accounted for subject specific information and gave accurate estimations for a narrow range of conditions [12,13,14]. Biomechanical simulations offer promise for energy expenditure estimation, but require detailed information such as marker data, accurate geometry, and properties of the subject’s musculoskeletal system [15, 16, 17, 18].

Data-driven methods used wearable sensors to explore portable energy expenditure estimation. Many different sensors were employed, such as accelerometers [19,20,21], activity monitors [22,23], heart rate monitors [24,25,26], electromyography (EMG) systems [27,28,29], and various combinations [30,31,32]. With the exception of [33], these studies used linear regression and hand-designed features to fit sensor data to estimate energy expenditure. While fitting informs the degree to which features correlate to energy expenditure, it does not evaluate the accuracy of the estimated energy expenditure for activity conditions or subjects not included in the correlation process. Benchmarking performance for energy expenditure estimation would enable clinicians and researchers to select models that meet their required level of accuracy for specific rehabilitation or research tasks.

The goal of this project was to estimate energy expenditure for three use cases as well as order the magnitude of energy expenditure across conditions. These use cases emulate experiments where different amounts of subject specific data are available and indirect calorimetry would be used. The three use cases estimated energy expenditure during walking conditions for subjects with some subject specific training data available, conditions for a new subject not included in the training data, and new conditions tested on a new subject. The input features were restricted to those measurable with wearable sensors and the length of input data per estimate consisted of a single gait cycle or fixed four second interval to enable portable and rapid estimation. Linear regression and neural network models were trained and tested on two datasets: (1) steady state walking with an ankle exoskeleton and (2) unassisted walking with a variety of loads and inclines. We assessed the error in energy expenditure estimation as well as the ability to order the magnitude of energy expenditure across conditions to evaluate the potential of this method for selecting optimal assistance conditions. Models were also trained with subsets of the input features to investigate their relative importance.

## Methods

### Datasets

The first dataset had subjects walk with an ankle exoskeleton providing a variety of assistance profiles, referenced in this paper as assisted walking [34]. Eight subjects were tested (7 men and 1 woman; age = 25.1 *±* 5.1 yr; body mass = 77.5 *±* 5.6 kg; leg length = 0.89 *±* 0.03 m). The subjects walked on an instrumented split-belt treadmill (Bertec, Columbus, OH) at 1.25 ms^*-*1^ for 8 minutes with the exoskeleton on one leg. Ground reaction forces, metabolic, and EMG data were collected. The EMG system (Trigno Wireless System; Delsys, Boston, MA) targeted the medial and lateral aspects of the soleus, medial and lateral gastrocnemius, tibialis anterior, vastus medialis, biceps femoris, and rectus femoris on both legs. Metabolic metrics were recorded with wireless metabolics equipment (Oxycon Mobile; CareFusion, San Diego, CA). The exoskeleton applied 9 different assistance strategies, with varying amounts of work and torque. The ground reaction forces and EMG signals were recorded at 2000 Hz, and all signals were recorded for the last 3 minutes of each condition once steady state was reached.

The second dataset investigated changes in energy expenditure when walking under loaded and incline conditions, referenced as inclined loaded walking [28,35]. We used data from all subjects (9 men and 4 women; age = 33.7 *±* 9.0 yr; body mass = 68.8 *±* 11.5 kg) who completed both loaded and incline studies. Subjects walked on an instrumented split-belt treadmill (model TMO8I with incline; Bertec Corporation, Columbus, OH). Ground reaction forces and moments, metabolic, and EMG data were collected. The EMG system (DE-2.1; DelSys, Boston, MA) targeted the soleus, medial gastrocnemius, tibialis anterior, medial and lateral hamstrings, vastus medialis, vastus lateralis, and rectus femoris. Metabolic metrics were recorded (Quark b2; Cosmed, Italy). Each subject walked under four loading conditions where 0%, 10%, 20%, or 30% of their bodyweight was added with a weighted vest. During each trial, the incline was set to 0%, 5%, and 10% grades for 5 minutes each. Thus, 12 walking conditions were recorded for each subject. The forces and EMG signals were recorded at 2000 Hz for the final 30 seconds of each condition and a metabolic measurement was collected continuously.

Separate data-driven models were trained on each dataset due to the different input signals. In summary, the assisted walking data had 22 time series signals with ground reactions forces for each foot and EMG signals from 8 muscles on both legs. The inclined loaded walking data had 14 time series signals due to ground reaction forces for each foot and EMG signals corresponding to 8 muscles on one leg. The measured energy expenditure values for the assisted data had a minimum and maximum of 269 W and 421 W with an average of 343 W. The inclined loaded data had extremes of 183 W and 892 W with an average of 478 W.

### Model architectures

Multiple types of models were considered to allow for energy expenditure estimation per gait cycle or fixed time intervals to enable estimates with any sets of sensors. Linear regression and neural network models were compared for estimating energy expenditure per gait cycle. The neural networks varied in size from 3 to 4 layers and 300 to 1000 neurons per layer. Dropout and L2 regularization were added to all layers to avoid overfitting to training data [36]. Rectified linear units were used as the activation functions for all neurons except the final fully connected layer. These networks were trained with a mean absolute error (MAE) loss function. Percent error was computed by scaling the MAE by the actual energy expenditure for each condition, giving a measure of relative accuracy. The reported MAE is normalized by average subject mass, although the networks output estimates in Watts.

Fixed time interval estimates needed to account for shifts in the time series data with temporal models. A recurrent neural network was used. The long short-term memory variant can capture long range dependencies and nonlinear dynamics [37]. The model tested had two long short-term memory layers of size 64, followed by a fully connected layer. A mean-squared error loss function was used in training while evaluation used MAE for consistency. The data was downsampled by averaging across every 32 sensor measurements to keep the length of the input data short enough for the model to perform well in recalling prior information. A four second interval of data recorded at 2000 Hz was downsampled to an input length of 250.

### Data processing

The inputs to the model consisted of ground reaction forces and EMG signals and the measured energy expenditure value was computed using indirect calorimetry. The three directions of ground reaction forces cannot be measured with wearable sensors, but prior work used force sensing insoles to estimate the three directions accurately [38]. The energy expenditure was calculated in Watts by passing the recorded oxygen and carbon dioxide values from each breath into the Brockway equations [39]. The ground truth energy expenditure value for each condition was found by averaging over all breaths during the last two minutes of recorded data, once steady state motion was achieved.

The force and EMG signals were filtered following standard biomechanics approaches. Both sets of signals were passed through a 4th order Butterworth filter. The force data was passed through a 30 Hz low pass filter to eliminate high-frequency noise. The EMG signals were filtered with a 30 to 500 Hz bandpass filter, rectified, filtered with a final 6 Hz low pass filter, and normalized by the maximum signal for each muscle during the normal walking trial.

The force and EMG time series data were formatted to make each sample of input data a fixed size. The first formatting method segmented the input signals by gait cycle, using the ground reaction forces to select data between right heel strikes. For a single gait cycle of approximately one second this results in a large and variable number of features to be fed into the estimation model. This variable length was converted to a fixed size of input data by dividing each input feature into a fixed number of bins, which were individually averaged. The number of bins was experimentally selected to be 30. Splitting data by gait cycle requires sensors to measure foot force or acceleration [40]. Another formatting method segmented data by fixed time intervals for use with sets of sensors that could not split the data by gait cycle. Every four seconds of data was taken as one input.

### Data analysis

Three common use cases were used to evaluate model performance: “condition”, “subject”, and “novel”. The condition use case estimated energy expenditure during conditions for subjects with some subject specific training data available. Random conditions were removed from the training data, often referred to as being held out of the training data, and treated as a test set to evaluate performance. These held out conditions were not necessarily the same conditions across all subjects. The condition use case simulated a use case where some subject training data was available and new conditions were tested. This is common in research when the same subjects repeat multiple experiments and subject specific data is available from those prior experiments. The subject use case held out one entire subject to estimate all conditions for this new individual. This is often seen in clinical or research work when a new subject performs a standard set of activities that prior subjects have completed. The novel use case held out one subject and the same conditions across all subjects. The novel use case represents an ideal case where a model generalizes energy expenditure estimation for a new subject completing a new task, neither of which are available in the training data.

A portion of the dataset, the validation set, was removed from the training data to evaluate the performance in order to tune the parameters of the neural network models. The conditions use case held out approximately 10% of the total conditions from any subjects. The subjects use case held out three subjects from the inclined loaded data and two subjects from the assisted data. The novel use case held out two complete subjects and two conditions across all subjects. The parameter values with the best performance on the validation set were selected and the validation set was replaced into the training set, due to the small number of subjects available. Thus, all data was used to determine the model accuracy.

The averaged model performance was measured using cross-validation. The entire dataset was divided into a number of sections equal to the number of subjects. Each section was iteratively treated as the test set, with the remaining sections combined to become the training set. The estimation accuracy was averaged across all test sets. Each test set for the condition use case consisted of roughly 10% of the total conditions from any subjects. The subject use case treated each subject as an individual test set. The novel use case treated each subject and two random conditions removed from all subjects as one test set; only these two conditions removed from the training data were estimated for the subject in the test set.

Confusion matrices visualized the difference in ordering the magnitude of energy expenditure across conditions for the model estimates and experimental measurements. The ground truth energy expenditure value of all conditions were ordered by increasing value along the vertical axis. The horizontal axis displayed the ordering estimated by the neural network models. The value of each grid square was normalized by the total number of subjects, thus a value of 1 corresponded to a perfect match between the measured and estimated ordering for that condition.

## Results

Neural network models estimated energy expenditure during assisted walking and inclined loaded walking within 5% of the measured values for the best subjects (Fig. 1). These estimations occurred per gait cycle and estimated all conditions for a new subject not included in the training data.

**Figure 1:**
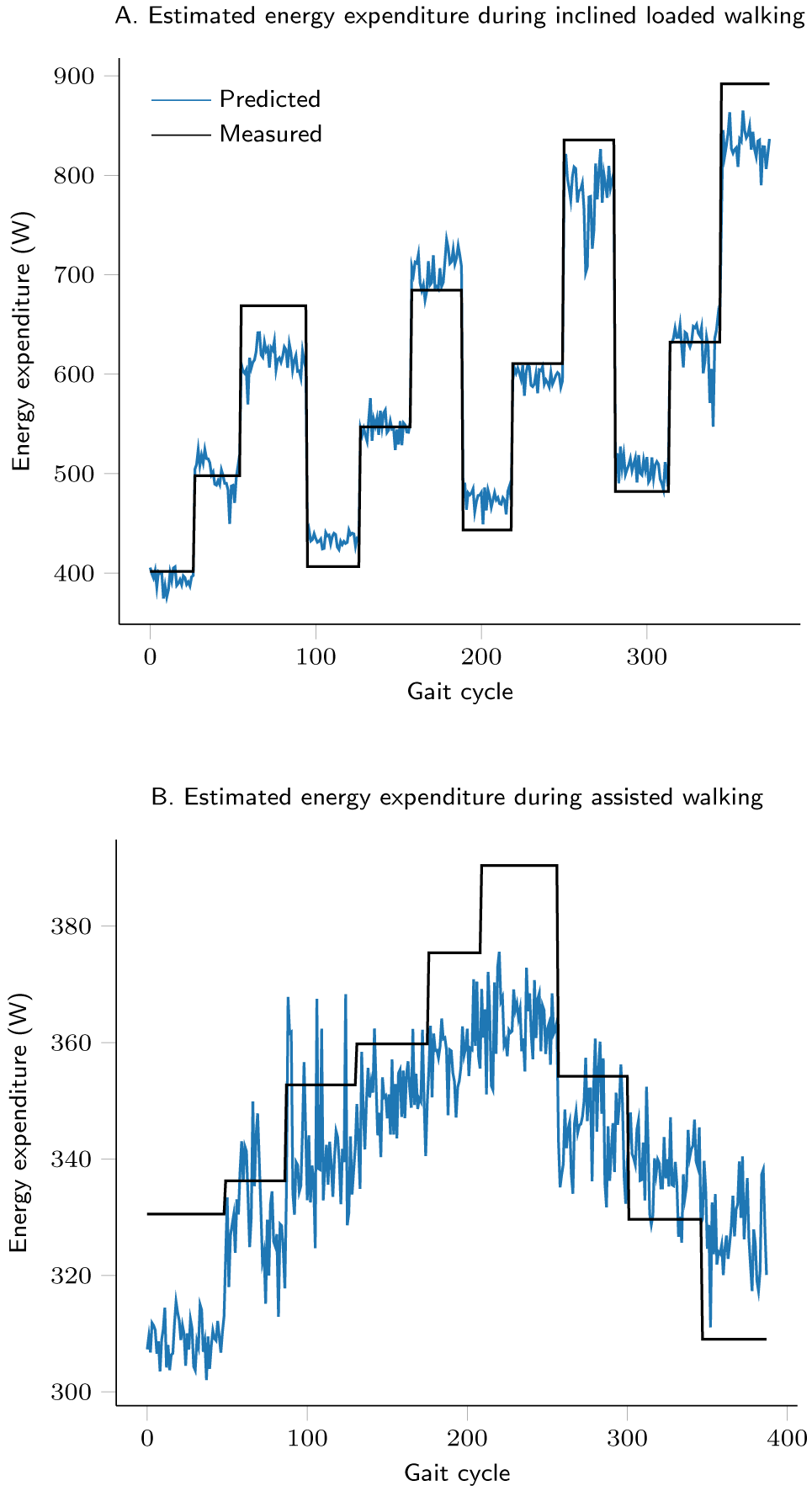
A visual comparison of neural network model estimates and measured energy expenditure values for the subject with lowest error when estimating all conditions in the dataset for a new subject.

The average error when using the neural network models to estimate energy expenditure during assisted walking for the three use cases was 4.4% for subjects with some subject specific training data available, 8.0% for conditions for a new subject not included in the training data, and 8.1% for new conditions tested on a new subject (Table 1). The linear regression models performed similarly to the neural network models. The average errors for the neural network models estimating energy expenditure during inclined loaded walking was approximately 2% worse in all use cases than during assisted walking (Table 2). The linear regression models performed only slightly worse, with an increase in the percent error between 0.6% and 2.4% compared to the neural network models. The R^2^ values for the linear regression models in all use cases during assisted and inclined loaded walking were greater than or equal to 0.96 when fitting training data.

**Table 1:** Comparison of linear regression and neural network energy expenditure estimates made per gait cycle for different use cases during assisted walking

| Model | Metric | Condition<sup>[a]</sup> | Subject<sup>[b]</sup> | Novel<sup>[c]</sup> |
| --- | --- | --- | --- | --- |
| Linear Regression | MAE (W/kg) | 0.18 | 0.37 | 0.37 |
|  | Error | 4.1% | 8.4% | 8.2% |
| Neural Network | MAE (W/kg) | 0.20 | 0.36 | 0.36 |
|  | Error | 4.4% | 8.0% | 8.1% |

<sup>[a]</sup>The condition use case randomly selected 10% of the conditions from any subjects as a test set, this was repeated as many times as there were subjects, with performance averaged across test sets.
<sup>[b]</sup>The subject use case removed one subject at a time from the training set to be the test set, averaging the performance across all subjects.
<sup>[c]</sup>The novel use case removed one subject at a time as well as two random conditions across all subjects from the training set. These removed conditions were estimated for the test set subject, with results averaged across all test sets.

**Table 2:** Comparison of linear regression and neural network energy expenditure estimates made per gait cycle for different use cases during inclined loaded walking

| Model | Metric | Condition | Subject | Novel | Subject Vertical Force<sup>[a]</sup> | Raw Subject<sup>[b]</sup> |
| --- | --- | --- | --- | --- | --- | --- |
| Linear Regression | MAE (W/kg) | 0.46 | 0.76 | 0.84 | 0.78 | 1.08 |
|  | Error | 6.7% | 12.1% | 13.7% | 12.3% | 16.5% |
| Neural Network | MAE (W/kg) | 0.43 | 0.61 | 0.70 | 0.63 | 0.70 |
|  | Error | 6.1% | 9.7 % | 11.7% | 10.0% | 11.2% |

<sup>[a]</sup>The subject vertical force use case was the subject use case with inputs restricted to vertical ground reaction forces and EMG signals
<sup>[b]</sup>The raw subject use case was the subject use case without any data preprocessing other than rectifying the EMG signals

A recurrent neural network using a fixed time interval of input data had an average error of 8.9% and MAE of 0.52 W/kg when estimating energy expenditure during all assisted walking conditions for a new subject. The recurrent neural network resulted in a 19.5% increase in MAE compared to the neural network estimating all conditions for a new subject per gait cycle.

The neural network models ordered the energy expenditure across all inclined and loaded walking conditions with a clear diagonal trend (Fig. 2A). The maximum and average percentage of correctly ordered conditions were 83% and 54%. The neural network ordering during assisted walking was less accurate, with additional errors causing a noisier diagonal trend (Fig. 2B). The recurrent neural network improved the clarity of the diagonal trend during assisted walking, but increased the spread of the outliers (Fig. 2C). The maximum and average percentage of correctly ordered conditions were 56% and 43%.

**Figure 2:**
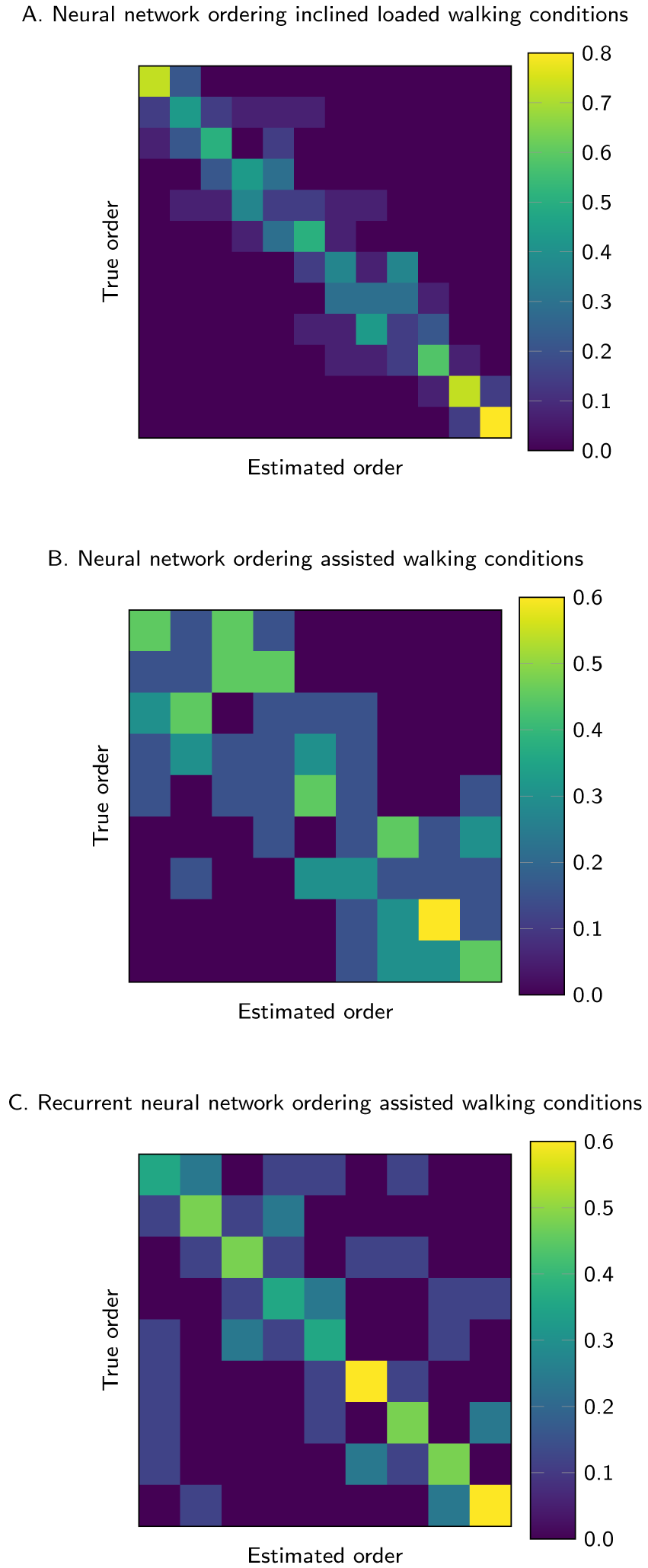
Visualized differences between the ordering of true, or measured, energy expenditure and the estimations across all conditions in the dataset for new subjects. The value in each grid square represents the number of estimated conditions ordered to match the corresponding true energy expenditure value. Perfect ordering results in a diagonal trend.

Restricting the input signals to EMG and the vertical ground reaction force increased the error by 0.3% compared to using all input signals (Table 2). Using all input signals but not performing any data processing other than rectifying the EMG signals increased the error for linear regression by 36% and neural networks by 15% compared to processed inputs.

Models restricting the inputs to either ground reaction forces or EMG achieved 8.1% or 9.2% error during assisted walking with a neural network (Table 3). Linear regression achieved similar levels of performance. For inclined loaded walking, the separated ground reaction force or EMG inputs achieved 11.5% or 23.9% error with a neural network and 12.5% or 31.6% error with linear regression. The addition of EMG to the linear regression model actually resulted in overfitting and degraded performance compared to only using ground reaction forces.

**Table 3:**
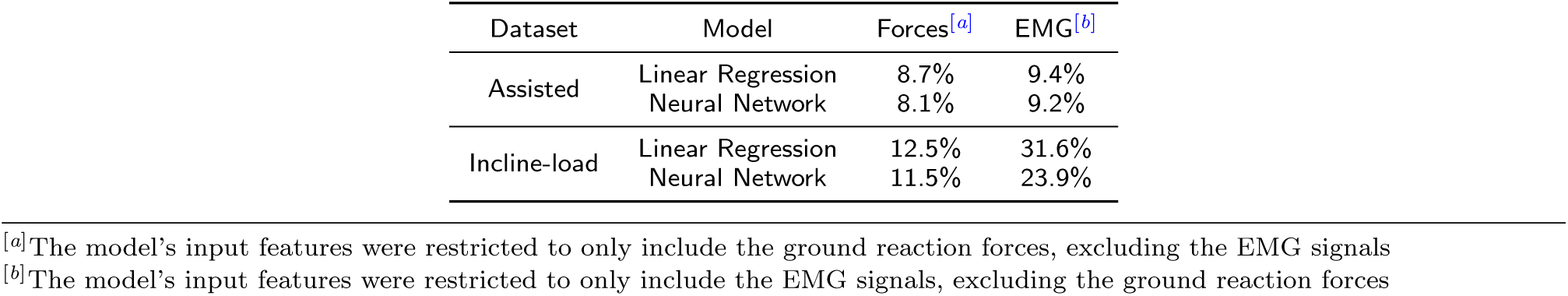
Average error for models with input features of either ground reaction forces or EMG signals when estimating energy expenditure during all conditions in the dataset for a new subject

| Dataset | Model | Forces<sup>[a]</sup> | EMG<sup>[b]</sup> |
| --- | --- | --- | --- |
| Assisted | Linear Regression | 8.7% | 9.4% |
|  | Neural Network | 8.1% | 9.2% |
| Incline-load | Linear Regression | 12.5% | 31.6% |
|  | Neural Network | 11.5% | 23.9% |

<sup>[a]</sup>The model's input features were restricted to only include the ground reaction forces, excluding the EMG signals
<sup>[b]</sup>The model's input features were restricted to only include the EMG signals, excluding the ground reaction forces

## Discussion

Understanding the computation the neural networks performs is challenging. Visualizations of the measured and estimated energy expenditure of the neural network model give insight into the model (Fig. 1). The larger range of energy expenditure values during inclined loaded walking indicates the ground reaction forces and EMG signals are more differentiated between conditions, which helped estimation consistency relative to assisted walking. The study that collected the assisted walking data noted two subjects were rejected for results more than two standard deviations from the mean [34]. The data for the rejected subjects were not available and not included in this work. This variability and small number of subjects provides some explanation for the noisy estimates.

When estimating energy expenditure for the use case where subjects had some subject specific training data available, the MAE and percent error was approximately halved compared to models estimating for conditions for a new subject not included in the training data (Tables 1 and 2). The similar performance between estimating energy expenditure for conditions for a new subject and new conditions for a new subject indicated the models captured the relationship between the inputs and energy expenditure, rather than just fitting a specific set of conditions found in the training data. The additional trainable weights in neural networks marginally improved performance over linear regression. The worse performance across all inclined loaded walking models was likely due to the larger range of energy expenditure values across conditions than during assisted walking (Table 2). The high R^2^ values from fitting linear regression training data indicated that using EMG and ground reaction forces as inputs with the binning structure was informative for estimating energy expenditure.

Capturing the general trends in energy expenditure across conditions is important for distinguishing the order of effort for a subject. The large range of energy expenditure values across inclined loaded conditions made ordering simpler than the assisted conditions. Due to these distinct conditions, the confusion matrix for the inclined loaded data had a clear diagonal trend with a maximum of 83% of conditions ordered correctly (Fig. 2A). The average percentage of correctly ordered conditions was much lower at 54%, but the errors occurred near the correct ordering, making the errors explainable. The difficulty of ordering the conditions during assisted walking due to the small range of energy expenditure was visualized with a less defined diagonal trend (Fig. 2B). The estimated ordering was within a few conditions of the actual ordering, indicating reasonable estimations. The recurrent neural network model ordering during assisted walking improved the clarity of the diagonal trend, but with outliers further from the correct conditions. This reflects the difficulty of ordering with a fixed time interval of input data rather than using the gait cycle structure for organizing temporal information (Fig. 2C).

The minimal increase in error when restricting inputs to the vertical ground reaction force and EMG signals indicated wearable sensors measuring normal force, such as force sensing insoles, may be capable of collecting the important force information (Table 2). The similar performance of the neural networks trained on raw or processed inputs shows promise for handling noisier wearable sensor data with minimal preprocessing.

Models with the input features restricted to either ground reaction forces or EMG signals had similar weights for both sets of input features during assisted walking (Table 3). Inclined loaded walking models relied more on the ground reaction forces, as these forces likely encapsulated information such as subject weight, incline, and amount of added load. Thus, the importance of certain features is dependent on the walking conditions, with a wider range of features enabling more robust performance when generalizing to new conditions. When using only EMG signals as input features, the neural networks again handled noisy signals well and outperformed linear regression. The recurrent neural network with input data in fixed time intervals offered slightly worse performance compared to the per gait cycle estimation, but would enable flexibility for use with any sensors.

### Limitations

The data-driven models could likely be improved by fine tuning the features and including more wearable sensor measurements. The linear regression features consisted of input signals individually averaged into a fixed number of bins for each gait cycle. Using feature selection or hand designing additional features may improve performance of linear regression.

A comparison between the performance of our models and prior work was excluded as prior studies correlated sensor signals to measured energy expenditure and report measures such as R^2^, but do not evaluate performance for any use cases similar to what we present. Even studies that report some measure of performance, such as the error in fitting the training data, cannot be directly compared as the range of energy expenditure across conditions, size of dataset, and the use case impact performance.

The input features used in this study could be collected with wearable sensors, but these sensors may add noise which would likely reduce performance. Including other wearable sensors investigated in previous research may improve estimation and generalization.

The similarity between the conditions in the training set and test set impacts the performance. When applying these models to other datasets, the same level of performance is expected if the new dataset has a similar size. A truly generalizable model for energy expenditure estimation would require significantly more data across a wider range of conditions and subjects. Hand designed experiments showed that estimating conditions with energy expenditure levels on the extremes of the dataset were most difficult. For example, holding out the two conditions where the exoskeleton applied the most work resulted in roughly three times the error compared to holding out random conditions with a linear regression model for the novel use case. To use these models in practice, the range of conditions to be tested should be similar to the conditions in the training data.

## Conclusions

This work benchmarks the performance of models used to rapidly estimate energy expenditure for use cases common in clinical and rehabilitation settings. If the performance of a model described here meets the requirement of a particular study, researchers may use these models rather than indirect calorimetry. The similar accuracy in estimating energy expenditure during conditions present in the dataset for a new subject and new conditions for a new subject suggests that the models learned a relationship between the input features and the energy expenditure, rather than just fitting conditions with similar training data. The models were also able to order the energy expenditure across conditions which could enable selection of optimal assistance conditions. Restricting input features to signals possible to measure with mobile sensors allows for flexible and scalable deployment of the models. These models take steps towards truly generalizable energy expenditure estimation which would improve rehabilitation and mobility beyond a lab setting.

## Abbreviations

EMG: Electromyography;
MAE: Mean absolute error

## Availability of data and material

The datasets analyzed in this work, code to train the models, and settings files for all models presented are available in the energy expenditure estimation repository on GitHub [41].

## Author’s contributions

All authors conceived the research questions. PS and RT performed the statistical analysis and wrote the manuscript draft. All authors reviewed, edited, and approved the final version of the manuscript.

## Acknowledgements

The authors thank Amy Silder and Rachel Jackson for access to and discussions about their data.

## References

1. Lauer, E.A., Houtenville, A.J.: Annual Disability Statistics Compendium: 2016 (2016)

2. Torburn, L., Powers, C.M., Guiterrez, R., Perry, J.: Energy expenditure during ambulation in dysvascular and traumatic below-knee amputees: a comparison of five prosthetic feet. Journal of Rehabilitation Research and Development 32, 111–119 (1995)

3. Brouwer, B., Parvataneni, K., Olney, S.J.: A comparison of gait biomechanics and metabolic requirements of overground and treadmill walking in people with stroke. Clinical Biomechanics 24(9), 729–734 (2009)

4. Zhang, J., Fiers, P., Witte, K.A., Jackson, R.W., Poggensee, K.L., Atkeson, C.G., Collins, S.H.: Human-in-the-loop optimization of exoskeleton assistance during walking. Science 356(6344), 1280–1284 (2017)

5. Felt, W., Selinger, J.C., Donelan, M.J., Remy, D.C.: “Body-in-the-loop”: Optimizing device parameters using measures of instantaneous energetic cost. PLoS ONE 10(8) (2015)

6. Kim, M., Ding, Y., Malcolm, P., Speeckaert, J., Siviy, C.J., Walsh, C.J., Kuindersma, S.: Human-in-the-loop Bayesian optimization of wearable device parameters. PLoS ONE 12(9) (2017)

7. Ding, Y., Kim, M., Kuindersma, S., Walsh, C.J.: Human-in-the-loop optimization of hip assistance with a soft exosuit during walking. Science Robotics 3(15) (2018)

8. Holdy, K.E.: Monitoring energy metabolism with indirect calorimetry: Instruments, interpretation, and clinical application. Nutrition in Clinical Practice 19(5), 447–454 (2004)

9. Van der Walt, W., Wyndham, C.: An equation for prediction of energy expenditure of walking and running. Journal of Applied Physiology 34(5), 559–563 (1973)

10. Pandolf, K., Haisman, M., Goldman, R.: Metabolic energy expenditure and terrain coefficients for walking on snow. Ergonomics 19(6), 683–690 (1976)

11. Duggan, A., Haisman, M.: Prediction of the metabolic cost of walking with and without loads. Ergonomics 35(4), 417–426 (1992)

12. Donelan, J.M., Kram, R., Kuo, A.D.: Mechanical work for step-to-step transitions is a major determinant of the metabolic cost of human walking. Journal of Experimental Biology 205(23), 3717–3727 (2002)

13. Kuo, A.D.: Energetics of actively powered locomotion using the simplest walking model. Journal of Biomechanical Engineering 124(1), 113–120 (2002)

14. Faraji, S., Wu, A.R., Ijspeert, A.J.: A simple model of mechanical effects to estimate metabolic cost of human walking. Scientific Reports 8(1) (2018)

15. Umberger, B.R., Gerritsen, K.G., Martin, P.E.: A model of human muscle energy expenditure. Computer Methods in Biomechanics and Biomedical Engineering 6(2), 99–111 (2003)

16. Delp, S.L., Anderson, F.C., Arnold, A.S., Loan, P., Habib, A., John, C.T., Guendelman, E., Thelen, D.G.: Opensim: Open-source software to create and analyze dynamic simulations of movement. IEEE Transactions on Biomedical Engineering 54(11), 1940–1950 (2007)

17. Uchida, T.K., Seth, A., Pouya, S., Dembia, C.L., Hicks, J.L., Delp, S.L.: Simulating ideal assistive devices to reduce the metabolic cost of running. PLoS ONE 11(9) (2016)

18. Dembia, C.L., Silder, A., Uchida, T.K., Hicks, J.L., Delp, S.L.: Simulating ideal assistive devices to reduce the metabolic cost of walking with heavy loads. PloS ONE 12(7) (2017)

19. Montoye, H.J., Washburn, R., Servais, S., Ertl, A., Webster, J.G., Nagle, F.J.: Estimation of energy expenditure by a portable accelerometer. Medicine and Science in Sports and Exercise 15(5), 403–407 (1983)

20. Swartz, A.M., Strath, S.J., Bassett, D.R., O’brien, W.L., King, G.A., Ainsworth, B.E.: Estimation of energy expenditure using CSA accelerometers at hip and wrist sites. Medicine & Science in Sports & Exercise 32(9), 450–456 (2000)

21. Crouter, S.E., Churilla, J.R., Bassett, D.R.: Estimating energy expenditure using accelerometers. European Journal of Applied Physiology 98(6), 601–612 (2006)

22. Heil, D.P.: Predicting activity energy expenditure using the Actical activity monitor. Research Quarterly for Exercise and Sport 77(1), 64–80 (2006)

23. Shcherbina, A., Mattsson, C.M., Waggott, D., Salisbury, H., Christle, J.W., Hastie, T., Wheeler, M.T., Ashley, E.A.: Accuracy in wrist-worn, sensor-based measurements of heart rate and energy expenditure in a diverse cohort. Journal of Personalized Medicine 7(2), 3 (2017)

24. Ceesay, S.M., Prentice, A.M., Day, K.C., Murgatroyd, P.R., Goldberg, G.R., Scott, W., Spurr, G.: The use of heart rate monitoring in the estimation of energy expenditure: a validation study using indirect whole-body calorimetry. British Journal of Nutrition 61(2), 175–186 (1989)

25. Brage, S., Brage, N., Franks, P.W., Ekelund, U., Wong, M.-Y., Andersen, L.B., Froberg, K., Wareham, N.J.: Branched equation modeling of simultaneous accelerometry and heart rate monitoring improves estimate of directly measured physical activity energy expenditure. Journal of Applied Physiology 96(1), 343–351 (2004)

26. Montgomery, P.G., Green, D.J., Etxebarria, N., Pyne, D.B., Saunders, P.U., Minahan, C.L.: Validation of heart rate monitor-based predictions of oxygen uptake and energy expenditure. The Journal of Strength & Conditioning Research 23(5), 1489–1495 (2009)

27. Wakeling, J.M., Blake, O.M., Wong, I., Rana, M., Lee, S.S.: Movement mechanics as a determinate of muscle structure, recruitment and coordination. Philosophical Transactions of the Royal Society of London: Biological Sciences 366(1570), 1554–1564 (2011)

28. Silder, A., Besier, T., Delp, S.L.: Predicting the metabolic cost of incline walking from muscle activity and walking mechanics. Journal of Biomechanics 45(10), 1842–1849 (2012)

29. Blake, O.M., Wakeling, J.M.: Estimating changes in metabolic power from emg. Springerplus 2(1), 229 (2013)

30. Eston, R.G., Rowlands, A.V., Ingledew, D.K.: Validity of heart rate, pedometry, and accelerometry for predicting the energy cost of children’s activities. Journal of Applied Physiology 84(1), 362–371 (1998)

31. Brage, S., Brage, N., Franks, P., Ekelund, U., Wareham, N.: Reliability and validity of the combined heart rate and movement sensor actiheart. European Journal of Clinical Nutrition 59(4), 561–570 (2005)

32. Ingraham, K.A., Ferris, D.P., Remy, C.D.: Using wearable physiological sensors to predict energy expenditure. In: IEEE International Conference on Rehabilitation Robotics (ICORR), pp. 340–345 (2017)

33. Vyas, N., Farringdon, J., Andre, D., Stivoric, J.I.: Machine learning and sensor fusion for estimating continuous energy expenditure. AI Magazine 33(2), 55–66 (2012)

34. Jackson, R.W., Collins, S.H.: An experimental comparison of the relative benefits of work and torque assistance in ankle exoskeletons. Journal of Applied Physiology 119(5), 541–557 (2015)

35. Silder, A., Delp, S.L., Besier, T.: Men and women adopt similar walking mechanics and muscle activation patterns during load carriage. Journal of Biomechanics 46(14), 2522–2528 (2013)

36. Srivastava, N., Hinton, G., Krizhevsky, A., Sutskever, I., Salakhutdinov, R.: Dropout: A simple way to prevent neural networks from overfitting. The Journal of Machine Learning Research 15(1), 1929–1958 (2014)

37. Hochreiter, S., Schmidhuber, J.: Long short-term memory. Neural Computation 9(8), 1735–1780 (1997)

38. Sim, T., Kwon, H., Oh, S.E., Joo, S.-B., Choi, A., Heo, H.M., Kim, K., Mun, J.H.: Predicting complete ground reaction forces and moments during gait with insole plantar pressure information using a wavelet neural network. Journal of Biomechanical Engineering 137(9) (2015)

39. Brockway, J.: Derivation of formulae used to calculate energy expenditure in man. Human Nutrition Clinical Nutrition 41(6), 463–471 (1987)

40. Tao, W., Liu, T., Zheng, R., Feng, H.: Gait analysis using wearable sensors. Sensors 12(2), 2255–2283 (2012)

41. Slade, P., Troutman, R., Kochenderfer, M.J., Collins, S.H., Delp, S.L.: Energy expenditure estimation project repository. https://github.com/pslade2/EnergyExpenditureEstimation. [Accessed 19 July 2018]

